# Bioinformatic Methods and Bridging of Assay Results for Reliable Tumor Mutational Burden Assessment in Non-Small Cell Lung Cancer

**DOI:** 10.1101/626143

**Authors:** Han Chang, Ariella Sasson, Sujaya Srinivasan, Ryan Golhar, Danielle M. Greenawalt, William J. Geese, George Green, Kim Zerba, Stefan Kirov, Joseph Szustakowski

**Affiliations:** Translational Medicine, Bristol-Myers Squibb, Princeton, NJ, USA

## Abstract

**Introduction:** Tumor mutational burden (TMB) has emerged as a clinically relevant biomarker that may be associated with immune checkpoint inhibitor efficacy. Standardization of TMB measurement is essential for implementing diagnostic tools to guide treatment.

**Objective:** Here we describe the in-depth evaluation of bioinformatic TMB analysis by whole exome sequencing (WES) in formalin-fixed, paraffin-embedded samples from a phase 3 clinical trial.

**Methods:** In the CheckMate 026 clinical trial, TMB was retrospectively assessed in 312 patients with non-small cell lung cancer (58% of the intent-to-treat population) who received first-line nivolumab treatment or standard-of-care chemotherapy. We examined the sensitivity of TMB assessment to bioinformatic filtering methods and assessed concordance between TMB data derived by WES and the FoundationOne^®^ CDx assay.

**Results:** TMB scores comprising synonymous, indel, frameshift, and nonsense mutations (all mutations) were 3.1-fold higher than data including missense mutations only, but values were highly correlated (Spearman’s r = 0.99). Scores from CheckMate 026 samples including missense mutations only were similar to those generated from data in The Cancer Genome Atlas, but those including all mutations were generally higher. Using databases for germline subtraction (instead of matched controls) showed a trend for race-dependent increases in TMB scores. WES and FoundationOne CDx outputs were highly correlated (Spearman’s r = 0.90).

**Conclusions:** Parameter variation can impact TMB calculations, highlighting the need for standardization. Encouragingly, differences between assays could be accounted for by empirical calibration, suggesting that reliable TMB assessment across assays, platforms, and centers is achievable.

**Key Points:**

- Tumor mutational burden (TMB) is a clinically relevant biomarker for efficacy of immunotherapy in patients with cancer
- Variations in TMB assessment parameters can shift the final TMB value. Harmonization and standardization are important to the successful clinical implementation of TMB testing
- TMB values assessed by different methods are highly correlated. Harmonization of TMB testing in patients with cancer is therefore achievable

## 1 Introduction

### 1.1 Tumor Mutational Burden

The genome of cancer cells can acquire genetic alterations that differ from the germline of the host [1]. Somatic mutation rates can be affected by exposure to exogenous factors, such as UV or tobacco smoke [2–4], or by compounding genetic defects, such as DNA mismatch repair deficiency, microsatellite instability, or replicative DNA polymerase mutations [2, 3, 5–8]. Clonal expansion results in a “mutational burden” that is propagated as the tumor evolves [9, 10].

Tumor mutational burden (TMB) is a quantitative assessment of the number of somatic mutations in coding regions of the tumor genome. TMB varies from patient to patient and also across different types of tumors [6, 11]. The average TMB value tends to be higher in some cancers, such as lung and melanoma, compared with other tumor types [6, 11]. Assessment of TMB involves next-generation sequencing (NGS) of tumor samples using one of several available platforms, and may also involve sequencing of normal patient tissue for germline variants. Initial exploratory analyses of TMB in patients with cancer [12–15] were carried out by whole exome sequencing (WES). WES is a comprehensive research tool to assess genomic alterations across the entire coding region of the ~22,000 genes in the human genome, comprising 1–2% of the genome [16, 17]. As an alternative to WES, the use of targeted cancer gene panels for TMB assessment is increasing [3, 4, 18–20]. The 324-gene panel assay FoundationOne^®^ CDx was recently granted premarket approval by the US Food and Drug Administration (FDA) for profiling of actionable mutations in solid tumors, and also provides assessment of genomic signatures such as TMB and microsatellite instability [21]. The FDA-authorized 468-gene panel assay MSK-IMPACT™ also captures TMB as part of its enhanced report for investigational use [4, 22], and Illumina’s TruSight Oncology 500 assay, which also captures TMB, has recently been granted Breakthrough Device Designation by the FDA [23]. Whereas TMB assessed by WES is typically reported as the total number of mutations per tumor, TMB outputs from gene panel assays are usually normalized to mutations per megabase (mut/Mb) because they differ in the number of genes and target region size [3, 4, 24, 25]. The precise calculation of TMB can, however, vary depending on the region of tumor genome sequenced, types of mutations included, and methods of subtracting germline variants [3, 26].

### 1.2 TMB and Immuno-oncology

Somatic mutations in coding regions of the genome (notably nonsynonymous mutations and, more significantly, frameshifts) potentially result in new or fragmented proteins/peptides (neoantigens), which can be recognized as “nonself” and elicit an antitumor immune response [1, 5, 26]. Neoantigens are therefore hypothesized to increase the immune cell repertoire and enhance the clinical efficacy of immune checkpoint inhibitors, such as anti–programmed death-1 (PD-1), anti–programmed death ligand 1 (PD-L1), and anti– cytotoxic T-lymphocyte antigen 4 (CTLA-4) [5, 27].

A leading hypothesis for the clinical implementation of TMB as a biomarker for immunotherapy efficacy is that high TMB is associated with increased antitumor immune responses and increased clinical benefit to immune checkpoint inhibitors, because TMB positively correlates with neoantigen load [5, 6, 13, 27–29]. This hypothesis is supported by the results of a number of clinical trials in several tumor types [12–14, 29–33]. In addition, a number of studies have reported that TMB prevalence and its association with immune checkpoint inhibitor efficacy are both independent of PD-L1 expression [12, 30, 33–36].

Several key studies of patients with non-small cell lung cancer (NSCLC) have demonstrated an association between high TMB and enhanced clinical benefit following first-line treatment with nivolumab (anti–PD-1) monotherapy and combination therapy. In the CheckMate 026 clinical trial (NCT02041533), patients with TMB in the upper tertile (≥243 missense mutations, as measured by WES) receiving nivolumab monotherapy showed an increased objective response rate (ORR) for nivolumab and longer median progression-free survival (PFS) compared with patients with TMB in the lower two tertiles (0–100 and 100–242 missense mutations, respectively) and with patients receiving chemotherapy [12]. CheckMate 568 (NCT02659059) used the FoundationOne CDx assay to establish a TMB cutoff (≥10 mut/Mb) for increased ORR with nivolumab + ipilimumab (anti–CTLA-4) combination therapy [32]; a cutoff of ≥10 mut/Mb was then chosen to define the “high TMB” patient population in CheckMate 227 (NCT02477826), which met its coprimary endpoint of PFS for nivolumab + ipilimumab vs chemotherapy in patients with high TMB, regardless of PD-L1 expression [30]. In patients with PD-L1 expression <1% who received nivolumab + chemotherapy, PFS was also longer in patients with high (vs low) TMB [37]. Thus, the CheckMate 227 trial validated the association of TMB with enhanced ORR and PFS with immunotherapy. The relationship between TMB and overall survival (OS) with immunotherapy, however, is less clearly defined. CheckMate 026 showed similar OS with nivolumab or chemotherapy, regardless of TMB status [12], whereas exploratory analysis of CheckMate 227 data showed a trend for prolonged OS with nivolumab + ipilimumab vs chemotherapy in the high and low TMB subgroups [38].

Other NSCLC studies showing enhanced benefit to immunotherapy in patients with high TMB used alternative WES methods or gene panel assays for TMB assessment from tumor or blood biopsies [29, 36, 39]. Such methods can differ in several factors, such as gene number, sequencing platform, sequencing depth, types of mutations that are included (e.g., nonsynonymous and/or synonymous single nucleotide variants [SNVs], short insertions/deletions [indels]), germline variant filtering methods, and format of test output (e.g., total mutations vs mut/Mb) [3, 4, 12–15, 24, 29, 36, 40–42]. Furthermore, patient groups can be selected around median, tertile, quartile, or fixed numerical values per tumor/per Mb [12–14, 29, 30, 35]. This makes interpretation of TMB assessment across different studies and platforms challenging. As the clinical utility of TMB assessment in tumors is established, treatment decisions may be made around a fixed TMB value, independent of the parameters that may influence this number. It is essential that the clinical community understands the meaning of TMB values generated by different methods and that guidance is provided for those establishing new approaches. Standardization of methods and clear definitions of assessment parameters are critical to the interpretation of TMB values into precision medicine for patients with cancer.

TMB assessment should accurately represent protein-altering tumor mutations that may contribute to enhanced antitumor immunity and thus improved responses to immune checkpoint inhibitors. In this study, we examine the analysis parameters and bioinformatics pipeline for reliable and accurate TMB assessment by WES, and highlight differences between this method and others. We illustrate the impact of including different types of mutations and germline filtering and explore the relationship between TMB assessed by WES and the gene panel assay FoundationOne CDx. In line with other harmonization efforts, the results highlight parameter variability across different approaches for TMB analysis and emphasize the importance of accurate reporting. Nevertheless, these results show that TMB data from different experimental platforms and bioinformatics pipelines are highly correlated, suggesting that assessment of TMB across different assays, platforms, and centers is achievable, and supporting the clinical implementation of TMB assessment for patients with cancer.

## 2 Methods

### 2.1 Samples for WES

TMB was retrospectively assessed by WES on formalin-fixed, paraffin-embedded (FFPE) tumor samples and matched blood from 312 patients with NSCLC (69 squamous and 243 nonsquamous), representing 58% of the intent-to-treat population in CheckMate026 [12]. Reasons for sample attrition included, but were not limited to, lack of patient pharmacogenetic consent, sample exhaustion by PD-L1 testing, and poor tissue sampling [12]. DNA and RNA were co-isolated from tumor tissue samples using the AllPrep DNA/RNA FFPE Kit (QIAGEN, Hilden, Germany) [12]. DNA from whole blood was isolated using the QIAamp DNA Blood Midi Kit (QIAGEN, Hilden, Germany) [12].

### 2.2 Library Preparation for WES

65–150 ng genomic DNA was fragmented to approximately 150 bp using the Covaris instrument (Covaris, Woburn, USA) and purified using Agencourt AMPure XP beads (Beckman Coulter, Indianapolis, USA). Libraries were prepared using the Agilent SureSelect^XT^ Reagent Kit (Agilent Technologies, Santa Clara, USA) with on-bead modifications [43]: the DNA was blunted and a single “A-tail” was added to each fragment; truncated PE P5 and P7 adapters were then ligated to each DNA fragment and the fragments were purified with AMPure XP beads. The DNA fragments were then amplified by polymerase chain reaction (PCR; eight cycles). Up to 500 ng of enriched library was used in the hybridization and captured with the SureSelect All Exon v5 bait (Agilent Technologies, Santa Clara, USA), enriching for 357,999 exons of 21,522 genes over a total target region of ~50 Mb [44].

Following hybridization, the captured libraries were purified according to the manufacturer’s recommendations and amplified by PCR (11 cycles) using a universal primer and a unique index primer specific to each library. This allowed molecules from each sample’s library to be distinguished from those of another sequencing library when several samples were processed simultaneously. As a result, multiple libraries could be combined or pooled prior to subsequent steps. The amplified product was checked for quality using the TapeStation (Agilent Technologies, Santa Clara, USA) and quantified by qPCR (Kapa Biosystems, Wilmington, USA). Normalized libraries were pooled and DNA was sequenced on the Illumina HiSeq 2500 using 2 × 100 bp paired-end reads.

### 2.3 WES Variant Calling Pipeline

A bioinformatic pipeline was used to filter NGS output data for germline variants, and detect and annotate synonymous, nonsynonymous (missense, nonsense, and frameshift) mutations and indels. A TMB score defined as total somatic mutations per tumor was derived (**Fig. 1**).

**Fig. 1.**
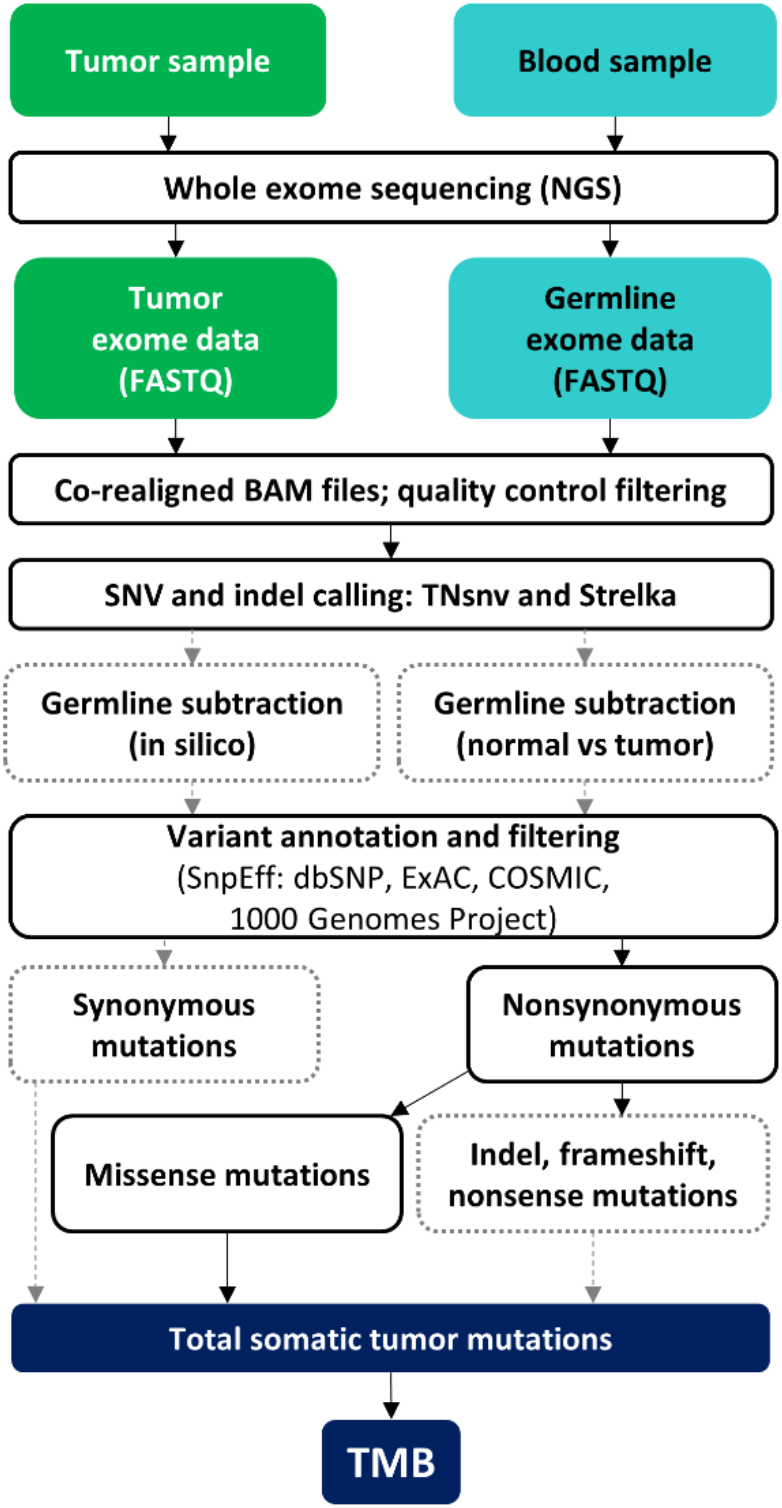
Workflow for TMB assessment by WES in this study. Solid black arrows denote steps that were included in all WES analyses. Dashed gray arrows show which steps were investigated for effects on TMB output in this study. COSMIC, Catalogue of Somatic Mutations in Cancer; ExAC, Exome Aggregation Consortium; indel, insertion/deletion; NGS, next-generation sequencing; SNV, single nucleotide variant; TMB, tumor mutational burden; WES, whole exome sequencing.

#### 2.3.1 Generation of BAM Files and Metrics from Raw FASTQ Reads

BAM files were generated from the paired FASTQ files following the Broad Institute’s best practices, using Sentieon Inc. implementation of the GATK pipeline [45]. The paired reads were aligned to the hg19 reference genome using the Burrows-Wheeler Aligner’s Maximal Exact Match (BWA-MEM) algorithm [46–48] and sorted; duplicate reads were marked. Indels were realigned and base quality scores recalibrated [49]. During this process, metrics were generated for total reads, aligned reads, and average coverage. Quality control filtering ensured that all samples used for analysis contained a total number of reads ≥45 million, mean target coverage ≥50×, and depth of coverage >20× at 80% of the targeted capture region or higher. If either tumor or blood data from a patient-matched pair failed any of these parameters, the pair was discarded [33].

The tumor and normal samples were processed individually as above to generate tumor and normal BAM files, which were then co-realigned. The BMS cohort-matcher tool (https://github.com/golharam/cohort-matcher), which utilizes BAM-matcher [50], compared the tumor and blood BAMs to ensure that they came from the same patient, in addition to checking for potential sample swaps within the cohort. If the genotype match between tumor and blood samples was <0.85, the pair was rejected from the final analysis.

#### 2.3.2 Variant Calling

The co-realigned (tumor + normal) BAM file, dbSNP [51], and target intervals consisting of coding exonic regions were used as the input for SNV calling and germline subtraction by the TNsnv somatic variant caller (Sentieon Inc., based on and mathematically identical to the Broad Institute’s MuTect) [52]. Default Sentieon TNsnv settings were used for analysis parameters that filter for sequence quality and variant allele frequency, including min_base_qual = 5, min_init_tumor_lod = 4, min_tumor_lod = 6.3, min_normal_lod = 2.2, contamination_frac = 0.02, min_cell_mutation_frac = 0, min_strand_bias_lod = 2 [53]. Somatic SNVs and indels were also called using the Strelka somatic variant caller using the tumor BAM file and normal BAM file for germline subtraction [54]. In Strelka’s BWA configuration file, the parameter “isSkipDepthFilters” was set to 1, as recommended for WES [46]. Three variant call format files (VCFs: one each for SNVs from TNsnv and Strelka, and a further VCF for indels from Strelka) were generated for each patient sample. To obtain somatic variants in the absence of a patient-matched normal sample, the tumor BAM and list of Catalogue of Somatic Mutations in Cancer (COSMIC) variants [55] were used as inputs for TNsnv, and HapMap NA12878 sequence data [56] were additionally used in place of a normal BAM in Strelka. VCFs were generated as above.

#### 2.3.3 Variant Annotation and Filtering

VCFs were filtered to retain only PASS variants. Annotations were then added using SnpEff, with RefSeq as the annotation source [57], from dbSNP [51], Exome Aggregation Consortium (ExAC) [58], COSMIC [55], and 1000 Genomes [59] databases. Any variants that were found in dbSNP, 1000 Genomes, and ExAC were excluded from the TMB calculation, unless they were also present in COSMIC. TMB was calculated as the total number of remaining mutations over a target region of ~30 Mb [60].

#### 2.3.4 Patient Characteristics

Patient characteristics were documented as part of the CheckMate 026 trial (NCT02041533). Race was self-reported by patients on enrollment, based on predefined categories.

### 2.4 Data Storage and Reprocessing

All necessary files and scripts required to reproduce the original results using the raw FASTQ files have been saved and versioned.

### 2.5 The Cancer Genome Atlas (TCGA) Data

WES data from CheckMate 026 samples were compared with data from 533 lung adenocarcinoma samples [61] and 177 squamous NSCLC samples [62] from the Broad Institute Firehose Genome Data Analysis Center (https://gdac.broadinstitute.org/). The results shown in this manuscript are based in part on data generated by the TCGA Research Network: http://cancergenome.nih.gov/.

### 2.6 FoundationOne CDx Assay

Investigational analysis of TMB by the FoundationOne CDx assay (https://www.foundationmedicine.com/genomic-testing/foundation-one-cdx. Accessed July 17, 2018) was carried out in Clinical Laboratory Improvement Amends (CLIA) certified laboratories at Foundation Medicine (Cambridge, MA, USA). NGS data was analyzed using proprietary software developed by Foundation Medicine [63] and quality control criteria that included tumor purity, DNA sample size, tissue sample size, library construction size, and hybrid capture yields were employed. Sequence data were mapped to the human genome (hg19) using Burrows-Wheeler Aligner (BWA) v0.5.9 [47], and PCR duplicate reads were removed and sequence metrics collected using Picard 1.47 (http://picard.sourceforge.net) and SAMtools 0.1.12a [64]. Local alignment optimization was performed using Genome Analysis Toolkit (GATK) 1.0.4705 [65]. Variant calling was performed only in genomic regions targeted by the test.

TMB was measured by counting all coding SNVs (synonymous and nonsynonymous) and indels present at ≥5% allele frequency and filtering out potential germline variants according to published databases of known germline polymorphisms, including dbSNP and ExAC. Additional germline variants still present after database querying were assessed for potential germline status and filtered out using a somatic-germline zygosity (SGZ) algorithm, a statistical model that considers tumor content, tumor ploidy and local copy number to account for rare germline variants [66]. Known and likely driver mutations were filtered out to exclude bias of the dataset. The resulting mutation number was then divided by the coding region corresponding to the number of total variants counted, or 793 kb, with the resulting number provided as mut/Mb [63].

In order to empirically calculate the conversion of TMB values between WES and the FoundationOne CDx assay, TMB results from the two assay methods were compared using nonparametric Passing–Bablok linear regression analysis that accounted for possible errors in both assays [67]. Following quality control filters on remaining CheckMate 026 samples for sample and data quality, which included tissue availability, tumor content, sequence target coverage, and sequencing depth, data from 44 samples were included in the comparison.

## 3 Results

### 3.1 Summary of TMB Effects on Clinical Outcome from CheckMate 026

In CheckMate 026, 541 patients with untreated stage IV or recurrent NSCLC and a PD-L1 tumor-expression level of ≥1% were randomly assigned to receive nivolumab or platinum-based chemotherapy [12]. WES libraries were successfully generated from 402 tumor samples and 452 blood samples. Three hundred twelve patient-matched tumor and blood samples passed quality control filters for TMB analysis (58% of the intent-to-treat population), with 158 of these patients receiving nivolumab and 154 receiving chemotherapy [12]. Patients were grouped into TMB tertiles based on their WES-derived somatic missense mutations (low [0–100], medium [100–242], high [≥243]). The results of the exploratory analysis published by Carbone et al. (2017) to test the hypothesis that patients with high TMB may derive enhanced benefit from nivolumab are summarized in **Table 1**. In patients receiving nivolumab, high TMB was associated with longer PFS compared with low or medium TMB, whereas high TMB was not associated with longer PFS in patients receiving chemotherapy [12].

**Table 1.** Summary of the effect of TMB on PFS in patients with NSCLC treated with nivolumab or chemotherapy

| TMB tertile<sup>a</sup> | Nivolumab (n = 158) |  | Chemotherapy (n = 154) |  | HR<sup>b</sup><br>(95% CI) |
| --- | --- | --- | --- | --- | --- |
|  | n | PFS, months<br>(95% CI) | n | PFS, months<br>(95% CI) |  |
| Low (0–100) | 62 | 4.2 (1.5–5.6) | 41 | 6.9 (5.4–NR) | 1.82<sup>c</sup><br>(1.3–2.6) |
| Medium<br>(100–242) | 49 | 3.6 (2.7–6.9) | 53 | 6.5 (4.3–8.6) |  |
| High (≥243) | 47 | 9.7 (5.1–NR) | 60 | 5.8 (4.2–8.5) | 0.62<br>(0.4–1.0) |

<sup>a</sup>Data adapted from Carbone et al [12]. TMB was defined as the total number of missense mutations. <sup>b</sup>HR for disease progression or death. <sup>c</sup>HR shown for low or medium TMB. CI, confidence interval; HR, hazard ratio; NR, not reached; NSCLC, non-small cell lung cancer; PFS, progression-free survival; TMB, tumor mutational burden.

### 3.2 WES Pipeline, Methodology, and the Impact of Variant Filtering on TMB Assessment

For the 312 CheckMate 026 patient samples described above, WES generated an average of 84 million reads per tumor sample (average 84.6× the mean tumor target coverage), and an average of 89 million reads per germline sample (average 93× the mean germline target coverage).

The bioinformatic analysis pipeline used is shown in **Fig. 1**. Output data from NGS were quality controlled and filtered to obtain a TMB score representing the total number of synonymous, indel, frameshift, missense and nonsense mutations (“all mutations”) per tumor sample. The impact of filtering data for missense mutations only was assessed. TMB scores including “all mutations” were highly correlated with matched data filtered for missense mutations only (Spearman’s r = 0.99; **Fig. 2**). Overall, TMB scores derived from “all mutations” were 3.1-fold higher than matched data filtered for missense mutations only. The median number of missense mutations was 170 and the median number of “all mutations” was 540.

**Fig. 2.**
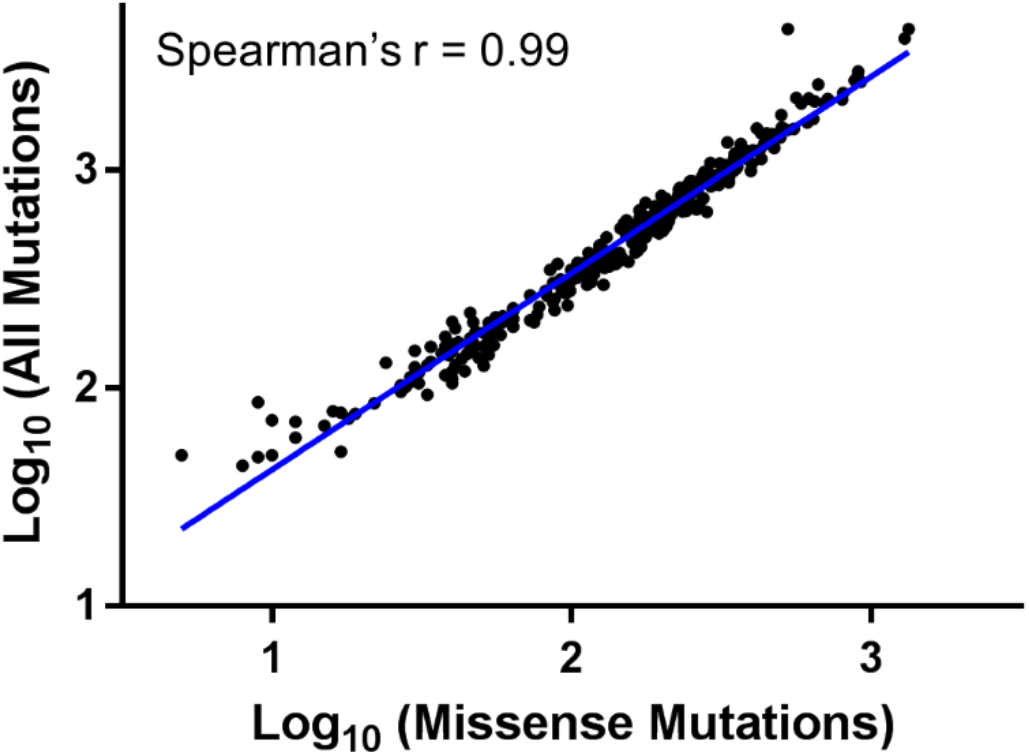
Analysis of TMB by WES in CheckMate 026 samples. Data show TMB scores including missense mutations only vs “all mutations” (including synonymous, indel, frameshift, missense and nonsense mutations). The linear regression line is shown in blue. TMB, tumor mutational burden; WES, whole exome sequencing.

The CheckMate 026 dataset was separated according to histology (69 squamous and 243 nonsquamous NSCLC samples) and compared with data from 177 squamous NSCLC and 533 lung adenocarcinoma (nonsquamous NSCLC) TCGA samples. For both histology types, TMB scores from CheckMate 026 were similar to those generated from TCGA when filtered for missense mutations only, with median values of 212 and 196 missense mutations observed for squamous samples, and 148 and 156 missense mutations observed for nonsquamous samples from CheckMate 026 and TCGA, respectively (**Fig. 3a**). When “all mutations” were included in the analysis, median TMB scores were higher across all groups (433 mutations in nonsquamous CheckMate 026 samples, 239 in nonsquamous TCGA samples, 704 in squamous CheckMate 026 samples, and 295 in squamous TCGA samples). Including “all mutations” in the TMB assessment had a larger effect on TMB scores from CheckMate 026 samples than for TCGA samples (**Fig. 3b**). A number of factors could have contributed to this variation, including differences in exome capture techniques or bioinformatic parameters used for variant calling. It is acknowledged, for example, that indel calling is more challenging than calling SNVs [68]. Investigators should be aware of the differences between various approaches to TMB assessment and ensure that technical details (eg, hybridization-capture libraries used and variant calling parameters) are sufficiently reported.

**Fig. 3.**
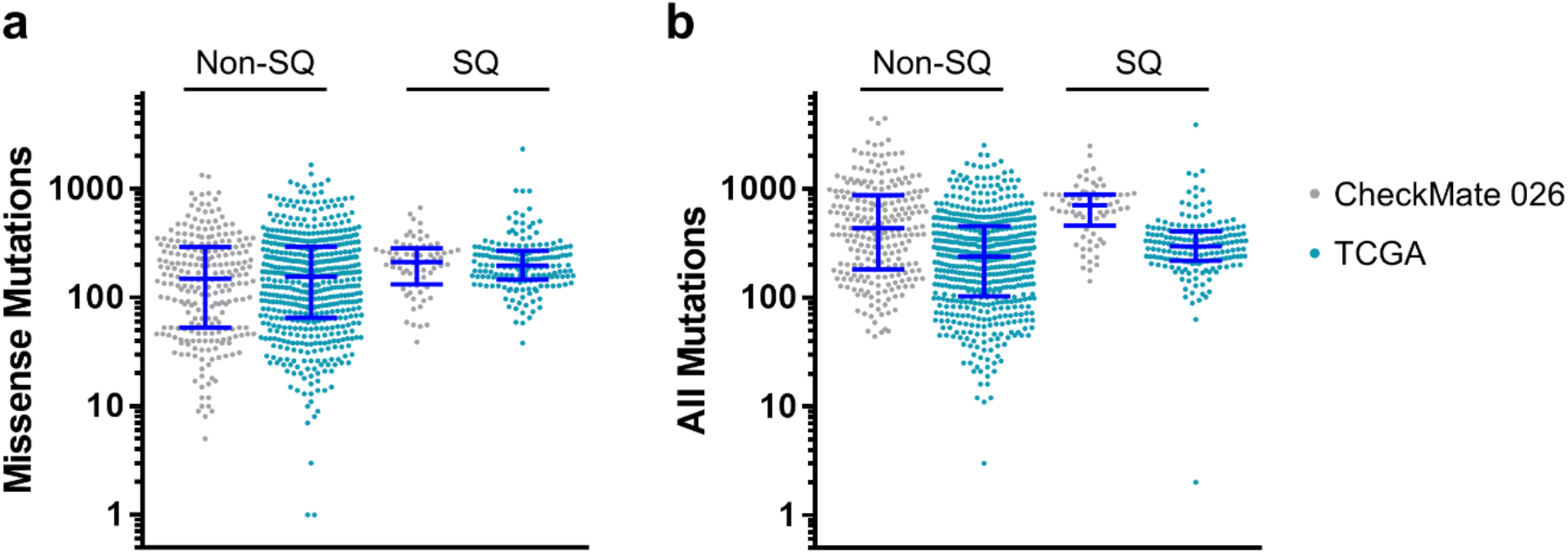
Comparison of TMB scores determined by WES in CheckMate 026 samples with those obtained from publicly available TCGA data. **(a)** TMB assessment includes missense mutations only. **(b)** TMB assessment includes “all mutations” (including synonymous, indel, frameshift, missense and nonsense mutations). Dark blue lines show the median value and interquartile ranges for each subset. Non-SQ, nonsquamous non-small cell lung cancer; SQ, squamous non-small cell lung cancer; TCGA, The Cancer Genome Atlas; TMB, tumor mutational burden; WES, whole exome sequencing.

### 3.3 Germline Filtering Using Tumor-Only and Patient-Matched WES Data

For all 312 NSCLC samples assessed for TMB in CheckMate 026, somatic mutations were identified using patient-matched blood (“normal”) samples to predict germline variants [12], with additional germline variants in the dbSNP, ExAC, and 1000 Genomes databases filtered out unless they were identified in COSMIC.

In a real-world setting, filtering germline variants using patient-matched “normal” samples may not be possible for ethical or practical reasons, and the tumor sample may be the only available sample for TMB testing. In silico methods to account for germline variants vary across different assays and may go beyond database filtering. To examine the significance of this potential limitation, TMB scores derived using patient-matched samples and database filtering (“tumor/normal TMB”) were compared with TMB scores derived solely from the tumor sample and in silico database filtering alone (“tumor-only TMB”). “Tumor-only TMB” values were on average approximately 100 mutations higher than “tumor/normal TMB” values (**Fig. 4a**). Data points fell noticeably into clusters with variable linear shifts in their intercepts. Grouping the dataset by patient characteristics revealed that the shifts in TMB value varied from 84 mutations in white patients to 132 mutations in black/African American patients and 213 mutations in Asian patients (**Fig. 4b**). These shifts were apparent throughout the whole dataset, including within data points around clinically relevant TMB cutoffs for medium or high TMB (100–300 mutations; **Fig. 4b**, inset).

**Fig. 4.**
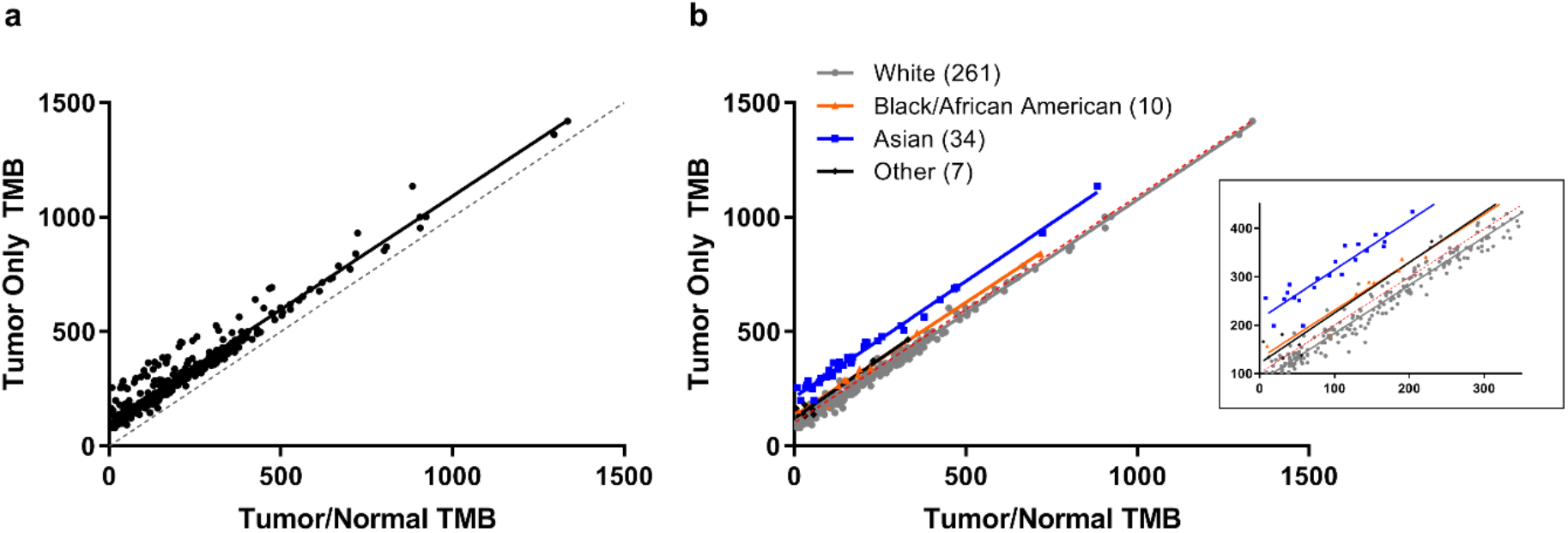
**(a)** The effect of germline subtraction by different methods on TMB values. TMB was derived from 312 patient samples by WES. The horizontal axis (tumor/normal TMB) shows TMB values calculated using patient-matched blood samples for germline subtraction. The vertical axis (tumor-only TMB) shows TMB values derived from the same sample, using public databases for in silico filtering. Black line shows linear regression across all patients. Equal values across the two datasets (X=Y) are represented by a gray dotted line. **(b)** Data from part (a) colored by patient race. Solid lines show linear regression analyses for subsets of patients grouped by race. Numbers in parentheses denote the sample numbers in each subgroup. Linear regression for the entire dataset is shown by a red dotted line. Inset shows the data magnified across tumor/normal TMB values from 0–350 mutations to highlight linear correlations occurring around TMB cutoff values that may be considered to be “clinically relevant”. TMB, tumor mutational burden; WES, whole exome sequencing.

The results above demonstrate that variations in analysis algorithms can offset the numerical output of the TMB assessment, but data from different pipelines are still highly correlated. This suggests that, as long as the specific parameters of the analysis pipeline (e.g., types of mutations included, databases used for filtering and annotation) are accurately reported, calibrations can be made using TMB data from different assays and data can be interpreted across various clinical studies.

While TMB assessments in the CheckMate 026 trial were calculated using matched normal blood samples for germline subtraction, those wishing to develop alternative TMB assessment methods should be aware that using publicly available data for estimating germline variation in place of matched normal samples may also influence TMB outputs, because current databases include uneven coverage of genetic variation across different races. This study therefore highlights that TMB assay methodology, variant calling parameters, and patient characteristics such as race should be considered when analyzing TMB, and that data from different methods may need to be adjusted in order to achieve comparable results and avoid potentially incorrect treatment decisions.

### 3.4 Bridging TMB Data between WES and the FoundationOne CDx Assay

We explored the relationship between TMB results generated by WES and the FoundationOne CDx assay in the CheckMate 026 samples in order to provide guidance to other investigators performing similar analyses and facilitate cross-trial comparisons. Variant allele determination by these two TMB assessment methods is dependent on a number of analytical differences, and simple arithmetic means are not sufficient to calibrate TMB values between them. An empirical method was therefore established to convert TMB values between WES and the FoundationOne CDx assay.

TMB was assessed in 44 FFPE samples from CheckMate 026 by the FoundationOne CDx assay, using the analysis pipeline shown in **Fig. 5**. Outputs from the FoundationOne CDx assay were compared by nonparametric linear regression with matched “tumor/normal TMB” data from WES. TMB values generated by these two assessment methods were highly correlated (Spearman’s r = 0.90; **Fig. 6a**). Consistent with concurrent studies that have established a clinically meaningful TMB cutoff in patients with NSCLC [30, 32], the FoundationOne CDx assay TMB cutoff level was evaluated at 10 mut/Mb. The precision of this TMB cutoff has been analytically validated, achieving >95% intra-run repeatability and >97% inter-run reproducibility [69]. TMB values generated by WES projected to an equivalent value of 199 missense mutations on WES (**Fig. 6a**). Patient samples were segregated according to whether they were < or ≥ the chosen cutoff values (**Fig. 6b**). Positive and negative agreements between WES and the FoundationOne CDx assay around the chosen values were 83% and 85%, respectively (95% confidence intervals [CIs] 61–94 and 67–94, respectively), and overall agreement between WES and the FoundationOne CDx assay was 84% (95% CI 71–92; **Table 2**).

**Fig. 5.**
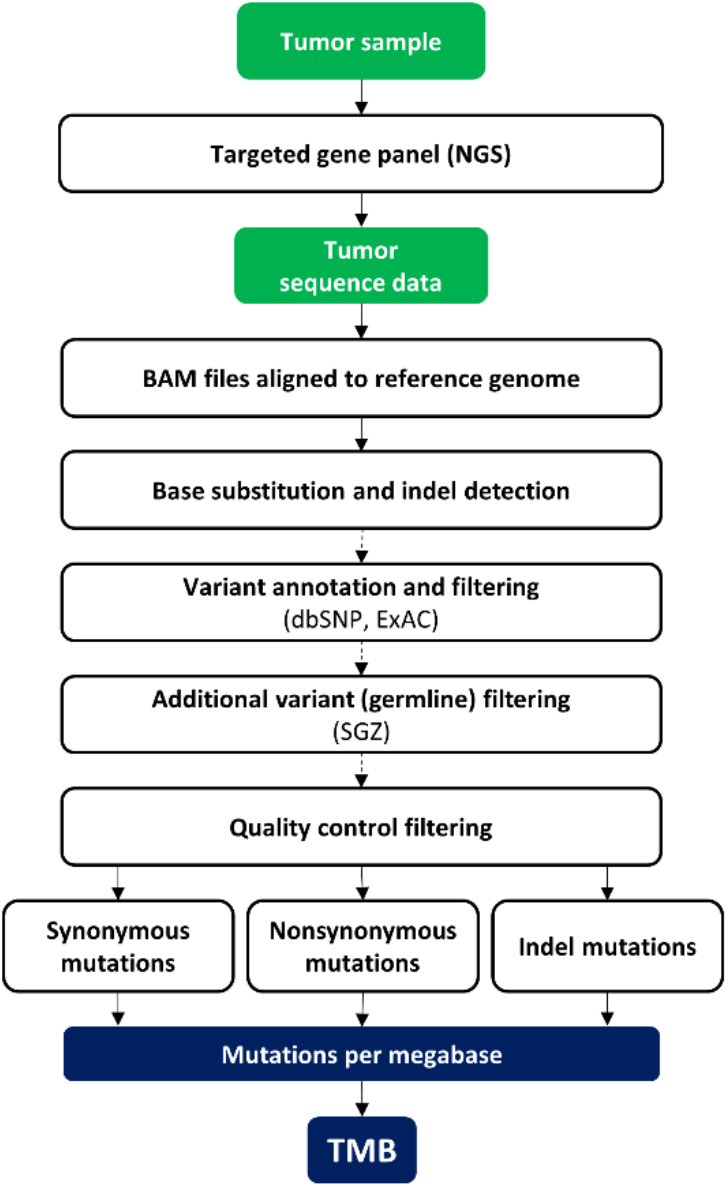
Workflow for TMB assessment using the FoundationOne CDx assay [63]. ExAC, Exome Aggregation Consortium; indel, insertion/deletion; NGS, next-generation sequencing; SGZ, somatic-germline zygosity; TMB, tumor mutational burden.

**Fig. 6.**
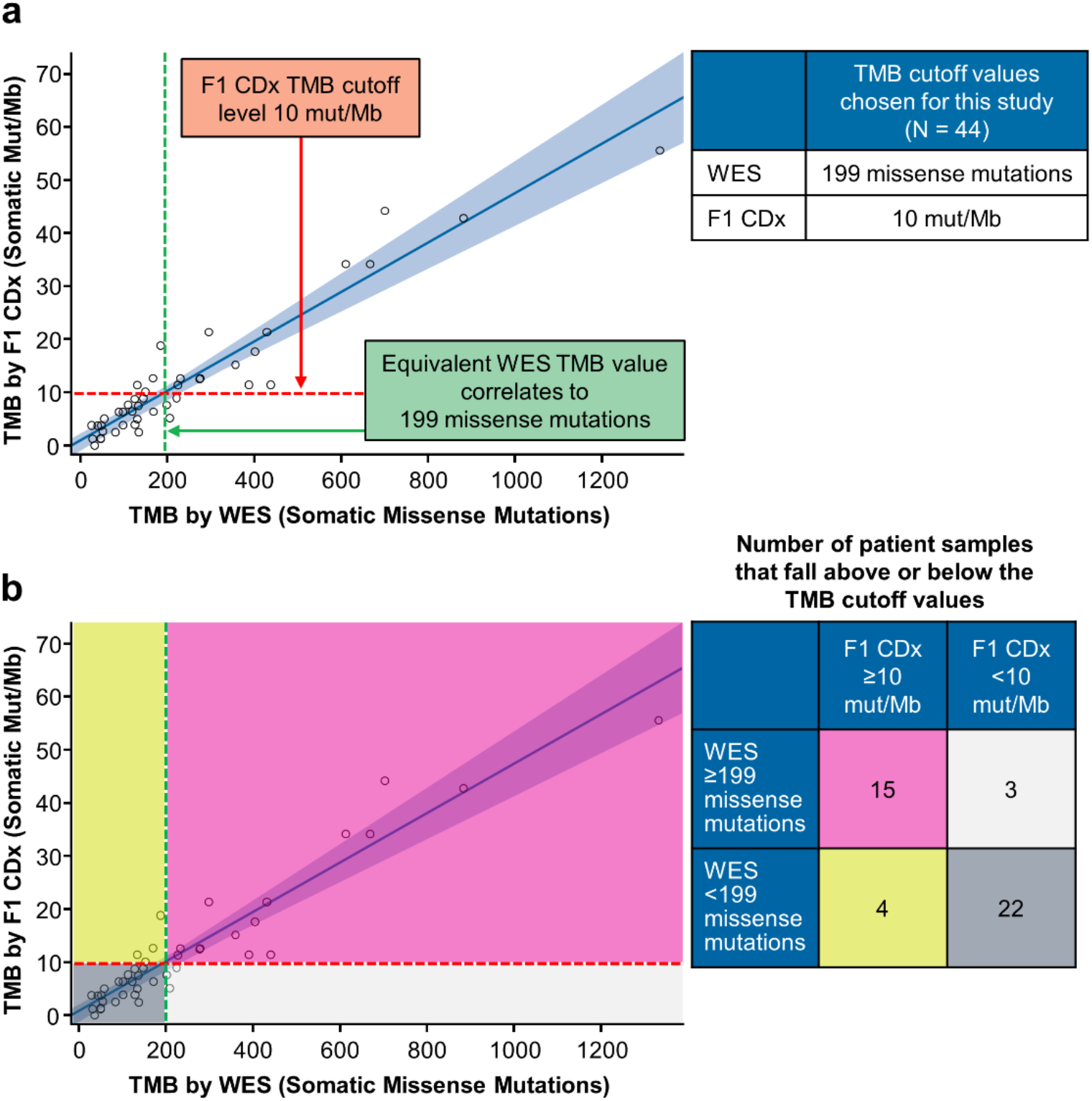
**(a)** Correlation of TMB assessed by WES and the FoundationOne CDx assay. Nonparametric linear regression is shown with a blue line. Blue shaded area shows 0.95– confidence bounds for the linear regression, calculated with a bootstrap (quantile) method. Cutoffs chosen for grouping of sample data are shown with red and green lines. **(b)** Grouping of TMB data into categories defined by cutoff values (10 mut/Mb by the FoundationOne CDx assay or 199 missense mutations by WES). F1 CDx, FoundationOne CDx; mut/Mb, mutations per megabase; TMB, tumor mutational burden; WES, whole exome sequencing.

**Table 2.** Agreement between TMB data derived by WES and the FoundationOne CDx assay

|  | Agreement (%) | 95% Wilson CI (%) |
| --- | --- | --- |
| Overall | 84 | 71–92 |
| Positive | 83 | 61–94 |
| Negative | 85 | 67–94 |

CI, confidence interval; TMB, tumor mutational burden; WES, whole exome sequencing.

## 4 Discussion

Response to therapy can be influenced by characteristics of the patient or tumor genome. Mutational burden within specific genes or pathways of interest is currently being investigated to facilitate the discovery of biomarkers for the clinical efficacy of a number of cancer treatments [2, 4, 18, 70, 71]. TMB represents the burden of mutations across the whole tumor exome, irrespective of gene function, acting as a surrogate biomarker for neoantigen load. Thus, high TMB may be a specific biomarker that is associated with improved efficacy of immunotherapy — a hypothesis that is supported by clinical evidence across several tumor types [5, 6, 12, 27–30, 32, 33, 37]. Evidence also suggests that TMB assessment may provide independent and complementary information to biomarker assessment by gene expression profiling or immunohistochemistry to guide treatment decisions [72–74].

A number of assays have been developed to determine TMB, with varying methodological parameters [24, 75]. In the initial analysis of TMB as a biomarker for immune checkpoint inhibitor efficacy, including CheckMate 026, TMB was retrospectively assessed by WES [12–15]. Targeted gene panels, however, are more readily interpretable and are considered to be a more pragmatic and potentially cost-effective approach to TMB testing in clinical diagnostics [3, 18, 24], and TMB was assessed by the FoundationOne CDx assay for the efficacy of nivolumab + ipilimumab and nivolumab + chemotherapy in CheckMate 568 and CheckMate 227 [30, 32, 37].

For successful real-world implementation of TMB testing, concordance between assay platforms used across different clinical trials should be established. Data from early studies using WES should therefore be converted to usable values for a clinical in vitro diagnostic, and vice versa. Because TMB represents the total quantity of somatic mutations across the tumor genome, several key attributes for quantifying TMB differ from those of NGS assays used to measure specific mutations. In the future, TMB will likely be assessed around specific cutoff values and patients with TMB above or below these values will be selected for different (individualized) treatment. Therefore, reliable, accurate, and reproducible real-world TMB testing options will be required across different clinical centers and countries, and analytical performance characteristics, such as precision and limit of quantification, play much larger roles in TMB assessment than in assays detecting presence or absence of targeted mutations [76–78].

Experiences during the implementation of PD-L1 testing in clinical practice as complementary or companion tests for immunotherapy have highlighted the need for standardization of reagents and methodologies, along with cross-center concordance studies [76–78]. It is therefore crucial and timely to take necessary measures toward the understanding and standardization of TMB assessment parameters, including different testing platforms, bioinformatic algorithms, and analysis workflow [79]. Systematic comparison and validation of new TMB assays with those used in clinical trials, harmonization of TMB data across alternative assays, thorough and accurate reporting, and empirical determination and calibration of cutoffs used to define patient populations will ensure the widespread availability of TMB testing. Worldwide efforts to ensure the harmonization of TMB assessment are ongoing [80, 81], including a European scheme from the International Quality Network for Pathology (IQN Path) [82], and a joint initiative between the US-based Friends of Cancer Research (Friends) and the Quality Assurance Initiative Pathology (QuIP) in Germany [75]. The Friends/QuIP partnership will use complementary in silico and laboratory-based approaches to introduce calibration standards and establish recommendations for reliable TMB assessment from WES and several different gene panel assays [75].

A number of factors vary between TMB assay methods, and it is important to be aware of these variations and ensure accurate reporting of each parameter. For example, target regions and databases used to call coding mutations can vary between different WES methods and from one gene panel assay to another [3, 4, 16, 17, 24, 60]. Sequencing depth also varies between assays. The average tumor sequencing depth for WES in the current study was 84.6×, whereas the FoundationOne CDx assay targets >500× median coverage with >99% of exons at >100× [63]. The FoundationOne CDx assay, like other gene panel assays, includes synonymous mutations in the TMB calculation to ensure a representative TMB estimation, whereas TMB assessed by WES may not [3, 4, 26]. The FoundationOne CDx assay involves a different sequence analysis pipeline and alternative software to that used for WES (see methods). Finally, to determine germline variants, WES incorporates a matched blood sample, whereas TMB scores obtained by the FoundationOne CDx are filtered for germline variants in silico using reference databases and the SGZ algorithm [3, 63, 66].

Consistent with published data, our results show that variations in assay parameters can cause a shift in the final TMB value [3, 4, 24, 29, 66]. For example, including synonymous, indel, frameshift, and nonsense mutations in the TMB calculation resulted in increased TMB levels compared with those including missense mutations only. Previous studies have also shown that race-dependent disparities can be mitigated using sufficiently comprehensive databases [42]. However, our findings suggest that shifts are still apparent after in silico filtering with dbSNP, ExAC, 1000 Genomes, and COSMIC, particularly in data from Asian patients. These results highlight the sensitivity of TMB assessment to in silico germline variant correction methods, and suggest that sophisticated bioinformatic adjustments beyond simple database filtering are required. The increased sequencing depth employed by existing assays such as FoundationOne CDx enables the precise measurement of allele frequencies. Furthermore, statistical models such as the SGZ algorithm use tumor-intrinsic factors to account for rare germline variants that may occur in smaller patient populations and be absent from public databases [63, 66].

Despite the large number of experimental differences between TMB assays, this study found that TMB values assessed by different methods correlated well, and calibration curves can be derived to convert TMB values between them. Patient-matched results that included “all mutations” vs “missense mutations” were highly correlated (Spearman’s r = 0.99). Furthermore, TMB values generated by WES and the FoundationOne CDx assay, which differ in methodology and bioinformatic methods, were also highly correlated (Spearman’s r = 0.90) and overall percentage agreement was 84% between these two assays (95% CI 71– 92). Further investigations and global standardization efforts are ongoing and include using reference standards to align experimental protocols and harmonize TMB assessment across different assay platforms [85, 86]. In conclusion, this study suggests that the clinical implementation of TMB testing is achievable and that educating and encouraging the clinical community to adopt accurate, reproducible assessment methods and reporting standards will ensure the streamlined implementation of TMB testing in patients with cancer.

## 5 Study Limitations

In this study, genomic analyses were carried out on FFPE samples. TMB assessment should consider the potential for sequencing artefacts introduced by formalin fixation. However, this method facilitates convenient sample storage and parallel biomarker testing by histology or immunohistochemistry. Thus, FFPE samples are the most commonly available samples for biomarker analyses [85].

While our methodological approach for TMB assessment by WES in CheckMate 026 samples involved the retrospective analysis of 312 samples, analysis of concordance between WES and FoundationOne CDx represents a pilot study using 44 samples. Increasing patient numbers is likely to improve the reliability of calibration across TMB assays and platforms. Furthermore, refining preanalytical procedures (e.g., sample fixation, microdissection, and DNA extraction), and optimization of analytical parameters (e.g., genome coverage, variant calling, and germline filtering methods) are likely to improve precision and reduce variation in TMB determination by gene panel assays.

In the WES analysis reported here, 21,522 genes were targeted over a total region of ~50 Mb and annotated by comparison with RefSeq over a region of ~30 Mb. The FoundationOne CDx assay targets ~1.8 Mb over 324 genes, of which ~0.8 Mb is used to calculate TMB [63]. Recent studies suggest that <0.5 Mb may be insufficient coverage for accurate TMB estimation when TMB is low and that panels covering >1 Mb achieve greater precision [3, 25, 83, 84]. The performance of gene panel assays should therefore be considered in the context of their intended use before selection.

## 6 Data Availability

Informed consent for data sharing was obtained from ~185 TMB-evaluable patients in the CheckMate 026 trial. Sequence data from these patients support the conclusions of this manuscript and have been deposited in the European Genome-phenome Archive (EGA), which is hosted by the European Bioinformatics Institute (EBI) and the Centre for Genomic Regulation (CRG), under accession number EGAS00001003661. *(Note for bioRxiv submission 2019-06-02: this data set will ‘go live’ in EGA mid-June 2019).*

## 7 Compliance with Ethical Standards

Results from the CheckMate 026 trial (ClinicalTrials.gov Identifier: NCT02041533) were reported previously [12]. The trial was conducted in accordance with the International Conference on Harmonisation Guidelines on Good Clinical Practice and the Declaration of Helsinki. Written informed consent was provided by all the patients before enrollment. HC, SS, AS, RG, DMG, SK, and JS designed the bioinformatics workflow for TMB analysis by WES and investigated the effects of parameter variation. HC, JS, GG, KZ, and WJG conceived, coordinated, and carried out the bridging study between WES and the FoundationOne CDx assay. JS and HC conceived the manuscript. All authors contributed to writing the manuscript and approved the final version. JS is the guarantor of this work and, as such, had full access to all of the data in the study and takes responsibility for the integrity of the data and the accuracy of the data analysis.

All authors are employed by, and own stock in, Bristol-Myers Squibb. In addition, HC, GG, WJG, and JS are inventors on one or more pending patent applications for tumor mutational burden as a predictive biomarker for immunotherapy.

## Acknowledgments

This work was funded by Bristol-Myers Squibb. We thank the patients and participating clinical study teams from CheckMate 026 who provided samples for this analysis, ONO Pharmaceutical Company Ltd. (Osaka, Japan) for collaborative development of the clinical trial program for nivolumab, and Foundation Medicine (Cambridge, MA, USA) for collaborative development of the FoundationOne CDx assay. Medical writing support and editorial assistance were provided by Stuart Rulten, PhD, and Jay Rathi, MA, of Spark Medica Inc, funded by Bristol-Myers Squibb, according to Good Publication Practice guidelines.

